# Rolling a mycobiome down a hill: endophytes in the Taiwanese Cloud Forest

**DOI:** 10.1101/210641

**Authors:** Dan Thomas, Roo Vandegrift, Yu-Ming Ju, Monica Hsieh, Bitty Roy

## Abstract

Fungal endophytes of plants are ubiquitous and important to host plant health. Despite their ecological importance, landscape-level patterns of microbial communities in plant hosts are not well-characterized. Fungal wood-inhabiting and foliar endophyte communities from multiple tree hosts were sampled at multiple spatial scales across a 25 ha subtropical research plot in northern Taiwan, using culture-free, community DNA amplicon sequencing methods. Fungal endophyte communities were distinct between leaves and wood, but the mycobiomes were highly variable across and within tree species. Of the variance that could be explained, host tree species was the most important driver of mycobiome community-composition. Within a single tree species, “core” mycobiomes were characterized using cooccurrence analysis. These core groups of endophytes in leaves and wood show divergent spatial patterns. For wood endophytes, a more consistent, “minimal” core mycobiome coexisted with the host across the extent of the study. For leaf endophytes, the core fungi resembled a more dynamic, “gradient” model of the core microbiome, changing across the topography and distance of the study.

## Introduction

Microbial community assembly and geographic patterns in microbes remain poorly understood, despite nearly a century of discussion (Baas-becking 1934 as cited in De Wit 2006, Martiny 2006, Green and Bohannan 2006, Peay 2010, Hanson 2012, Nemergut 2013). Host-associated microbes present additional complexity in modeling microbial community assembly, and raise questions concerning fidelity of host-microbe interactions. Rich microbial communities appear to be associated with all large, eukaryotic organisms (Rosenburg 2010, Hoffman 2010). Plant-fungal symbioses are important ( Malloch 1980, Stukenbrock 2008, Vandenkoornhuyse 2015) and at least as ancient as vascular plants (Redecker 2000, Krings 2007). Fungal endophytes, or fungi that live internally in plant tissues without incurring disease symptoms (Wilson 1995), have been shown to be widespread and important to plant health (Arnold 2003, Mejia 2008, Rodriguez 2009, Porras-Alfaro 2011). The endophytic compartment in which they reside is a distinct ecological space, in the sense that very different communities of microbes are observed outside vs. inside plant tissues (Santamaria 2005, Lundberg 2012, Bodenhausen 2013), at least partly due to host-microbe preferences (Schulz 1999, Oldroyd 2013, Venkateshwaran 2013). Plant organs have been shown to host distinct communities of endophytes (Bodenhausen 2013, Persoh 2013, Tateno 2014, Edwards 2015). Endophyte communities are also influenced by environmental conditions (Carroll 1978, Arnold 2003, Zimmerman 2012), in spite of presumed buffering from environmental stresses by host tissues. Fungal communities are subject to spatial processes such as dispersal limitation (Peay 2010, Higgins 2014). Fungal endophytes, therefore, make ideal systems for studying the interplay of host-microbe interactions, environmental influences, and spatial patterning of both host and microbes in natural settings.

The potential importance of microbes in adding ecological functions to their hosts (Rodriguez 2009, Johnson 2012, Woodward 2012) has led some to suggest that multicellular organisms may host *core microbiomes* (Hamady 2009, Shade 2011, Vandenkoornhuyse 2015), which are subsets of important and consistent microbial partners. Initial explorations of plant core microbiomes have been highly controlled (Lundberg 2012, Edwards 2015). Studies of plant-associated microbiomes in natural settings have rarely been framed in terms of core microbiomes (Kim 2011, Zimmerman 2012, Bodenhausen 2013, Higgins 2014, Kembel 2014). This is not a coincidence: outside of experimental settings, the prospect of detecting a cadre of microorganisms absolutely loyal to their host in the face of a complex and dynamic natural environment is daunting. This definition of the core microbiome, known as either a “substantial” or “minimal” core (Hamady 2009) may be useful when carefully applied to long-studied symbioses such as ruminant gut communities (Liggenstoffer 2010) or mycorrhizal relationships (Malloch 1980, van der Heijden 2009). This definition may not always serve for describing the numerous and labyrinthine microbe-host interactions that exist outside of laboratory settings. However, other definitions of core microbiomes exist that may be more useful for ecologically modeling microbiomes (Hamady 2009).

Here we acknowledge that plant hosts exert strong influence on community membership of their endophytic compartment. However, we hypothesized that even the most faithful fungal associates will uncouple from their hosts with changing environmental conditions and dispersal constraints. We predicted, on the scale of the present study, that plant mycobiomes resemble “gradient” core microbiomes (Hamady 2009). Under this model, microbiomes can totally change across a landscape, with host-interactions mitigating but ultimately not preventing environmentally- and spatially-driven changes in the microbiome. To test this, we compare community composition and ecological drivers between wood and leaf fungal endophytes in multiple species of plant host, to identify instances of differential response by microbial communities from host to environmental changes or spatial constraints. We map spatial patterns in the most strongly associated endophytic fungi of a single host species, to examine patterns of turnover in a putative core microbiome.

## Methods

Background/Site: Sampling occurred in summer of 2013 at Fushan forest, in Northeastern Taiwan (24º 45' 40” N, 121º 33' 28” E), which hosts a 25-ha Smithsonian-associated (Losos & Leigh 2004) Forest Dynamics Plot (FDP) (Su 2007). Fushan is a humid subtropical old-growth montane site that receives 4.27 m of rain each year. Most of this precipitation falls during rainy, cool winters, though a significant fraction of this rain is due to typhoons, the main agent of disturbance in this system, during warm summer months. The flora is diverse, characterized by many evergreen broadleaf tree species and a diverse understory of lianas, ferns, tree ferns, and other herbs, gramminoids, and shrubs. Vegetative communities can be broadly categorized into four community types described by dominant tree species combinations (Fig. 1). Topography is highly variable, with a maximum elevation of 733 m above sea level at an approximately central hilltop within the FDP, and a minimum of 600 m, though the present study sampled areas only as low as 650 m. The central hilltop adjoins lowland habitat with perennial streams along its eastern and southern bases, and mid-elevation upland habitat to the north. Perennial streams join and exit the FDP through a steep valley in the southwest of the plot (Fig. 2). The complex topography of Fushan has been summarized by classification of each 20 m x 20 m quadrat of the FDP into one of seven habitat types, based on aspect, slope, convexity, and elevation (Fig. 1), which are found to influence vegetative communities (Su 2010). Soil at Fushan FDP are generally acidic, with low fertility and organic carbon content. Soils are relatively young (inceptisols) due to erosion on steep slopes and flooding disturbances in lowland habitat. High leaching and erosion cause lower nutrient levels to occur in the central hilltop. See Su et al. (2007) for more details.

**Figure 1.**
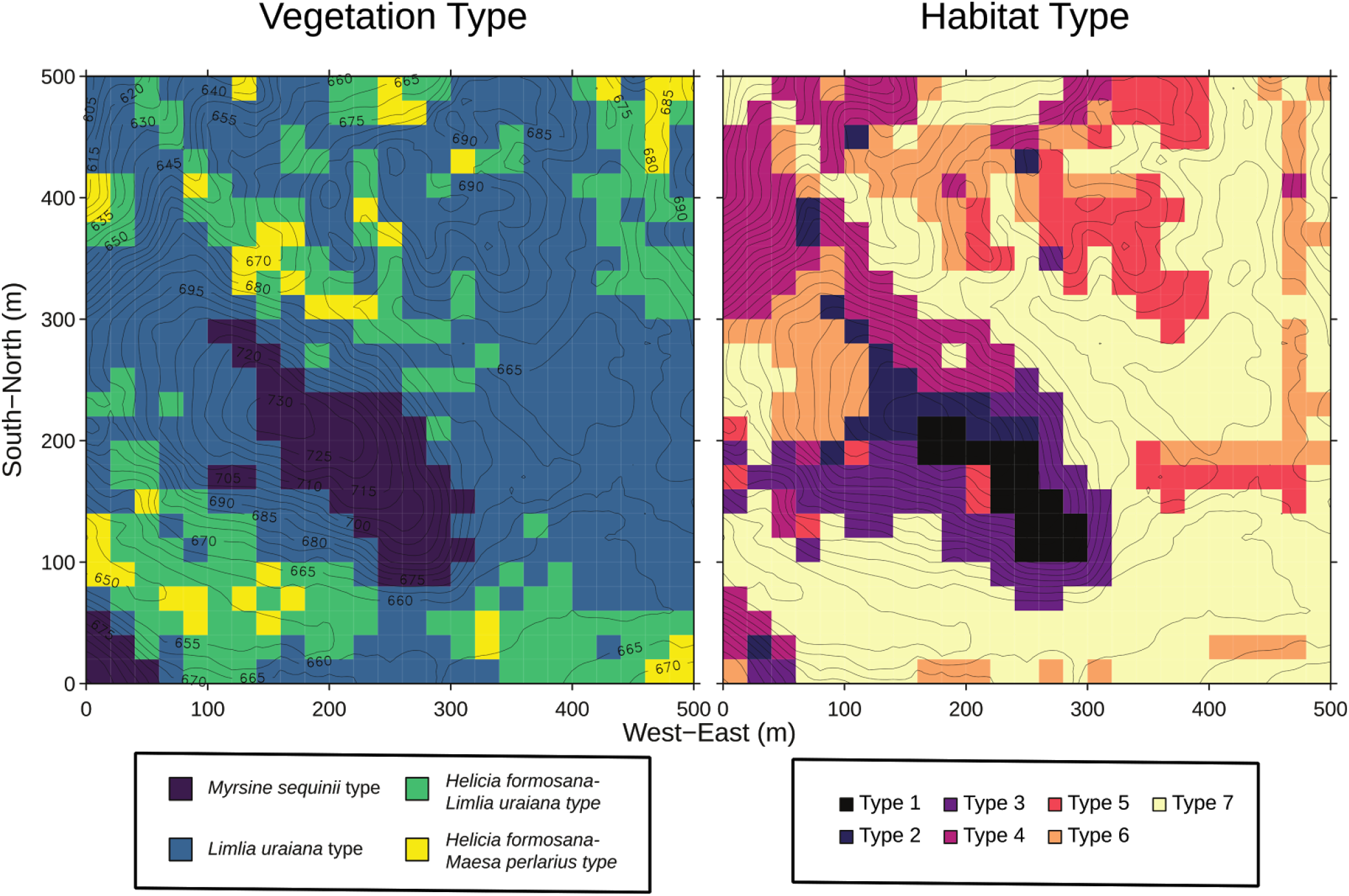
Left: topographic map of the Fushan FDP with the four vegetation types as classified by Su et al. ( 2007) Right: map of the habitat type, a composite classification based on microtopographic characteristics of quadrats, defined by Su et al. (2010). The units of the coordinates and contours are in meters, with quadrats at 20x20m scale. Figures reproduced with permission from authors. Click here for a higher resolution image.

**Figure 2.**
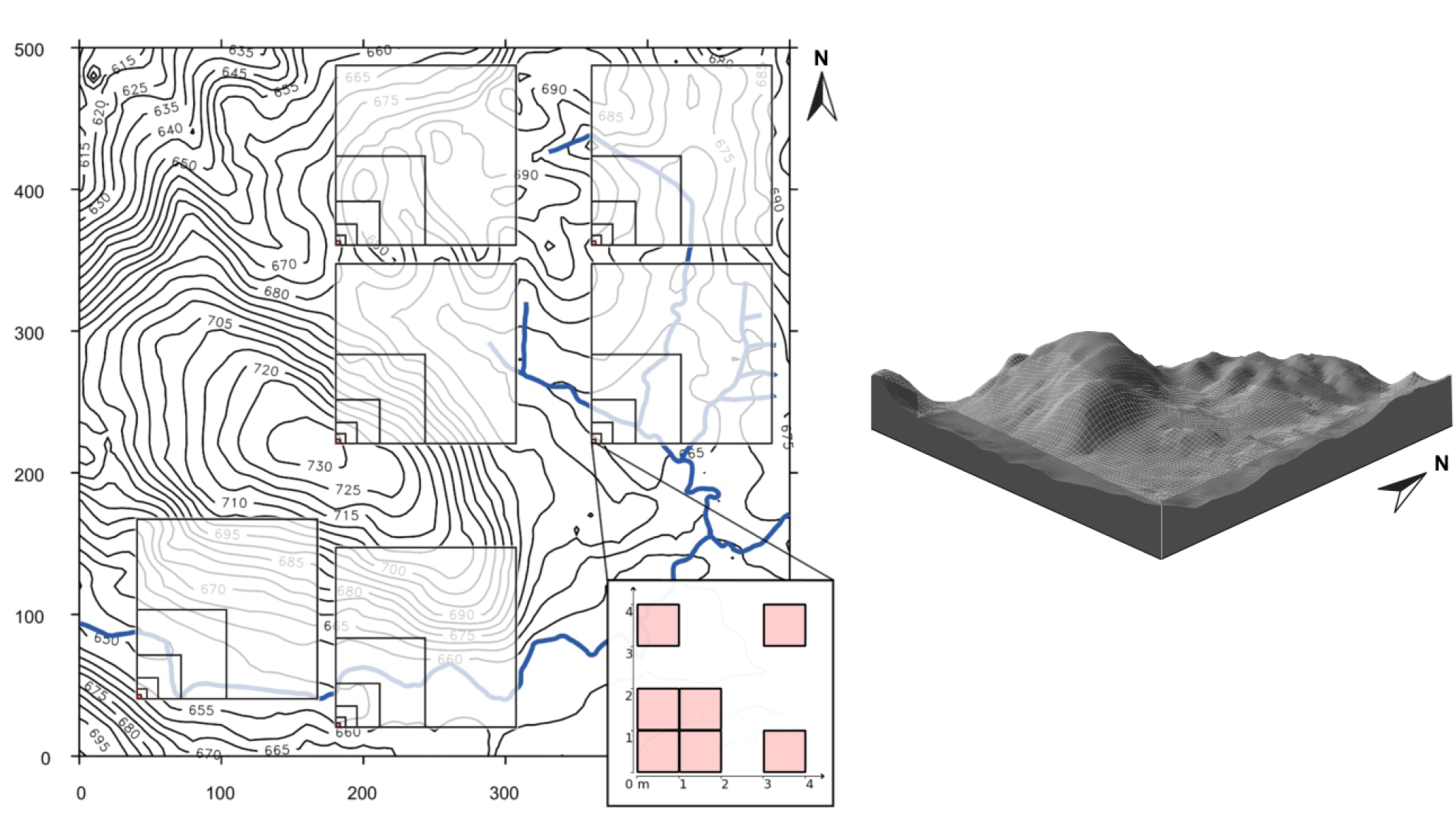
Left - An overview of nested-squares, logarithmic sampling scheme Vandegrift (2016). Vertices of squares are sample sites. Units are meters. Right - Perspective diagram of Fushan Forest Dynamics Plot (Su 2010). Figures reproduced with permission from authors. Click here for a higher resolution image.

## Field methods

Fushan FDP was divided into 9 sub-plots, and subplots were sampled using a nested logarithmic scheme intended to detect dispersal limitation and community turnover (Rodrigues 2013) (Fig. 2). Sampling of each set of nested points was undertaken in random order. Once sampling of a single set of nested squares had begun, all points within that set of nested points were sampled prior to beginning another. Six out of nine sets of nested squares were sampled, due to time constraints.

For each sampling point, we located the tree with the largest DBH with canopy above the point and collected the three lowest “healthy” appearing leaves that were safely reachable. Leaves and accompanying woody stems were obtained using a 3m collapsible pole pruner. Identification of host-tree was supplied by survey data from ongoing ecological research at Fushan FDP (Su 2007). All plant material was carried to a nearby field station and stored at 4°C for no longer than 5 days before processing.

## Lab methods

Preparation and sequencing of Illumina libraries for leaves and wood were undertaken separately, with differing protocols. Protocols for leaf fungal endophyte amplicon library preparations are given in Vandegrift (2016). Protocols for wood endophytes are given in detail in Thomas (2017). Briefly, all leaves were washed and surface-sterilized, and woody stem material was debarked with a sterile scalpel and phloem and sapwood were harvested. DNA was extracted from both in separate library preparations and ITS region 1 was amplified using a fungal-specific primer set with illumina© tagged, barcoded primers. Positive, “mock community” controls were included in the wood-endophyte library, and pure-water negative controls were included in both libraries. Samples were multiplexed and sequenced in separate illumina© Mi-Seq sequencer runs.

## Bioinformatics

Details of the bioinformatics pipeline are explained in Thomas (2017). Full scripts available in supplementary information (available here and here). Briefly, general bioinformatics protocols followed the USEARCH/UPARSE pipeline version 8.1 (Edgar 2013) wherever possible. Libraries of leaf and wood fungal endophyte DNA were prepared separately, so to maximize comparability, the reads from both libraries were combined as early as possible in the bioinformatics pipeline, following merging of paired ends. Variance stabilization of combined wood and leaf reads was done using using the *DESeq2* package in R (Love 2014, McMurdie 2013), using leaf/wood as the design variable. Positive controls were used to calibrate OTU similarity radius and minimum cutoffs, which were subtracted from all observations to reduce error from index-misassignment and artificial splitting of OTUs. Large differences in abundances remained among positive control OTUs even after variance stabilization, so all statistical analyses were conducted with incidence (presence/absence)-transformed community matrices.

## Statistical methods

### Overview

Ecological patterns of the entire fungal community of leaves and wood of all hosts were examined first. Analyses then were focused on patterns in the mycobiome of the single, most commonly-sampled host tree, *Helicia formosana*. Finally, host-fungus coccurrence patterns were used to define a core mycobiome that was also examined for ecological patterns (Fig. 3). Statistical analysis was conducted in R Statistical Software, version 3.3.1 (R core team 2016), with the *vegan* (Oksanen 2017), *phyloseq* (McMurdie 2013), *cooccur* (Griffith 2016), *igraph* (Csardi 2006) and *ecodist* (Goslee 2007) packages. Where required, all endophyte community comparisons were conducted using Bray-Curtis dissimilarity index (Bray 1957, McCune 2002).

**Figure 3.**
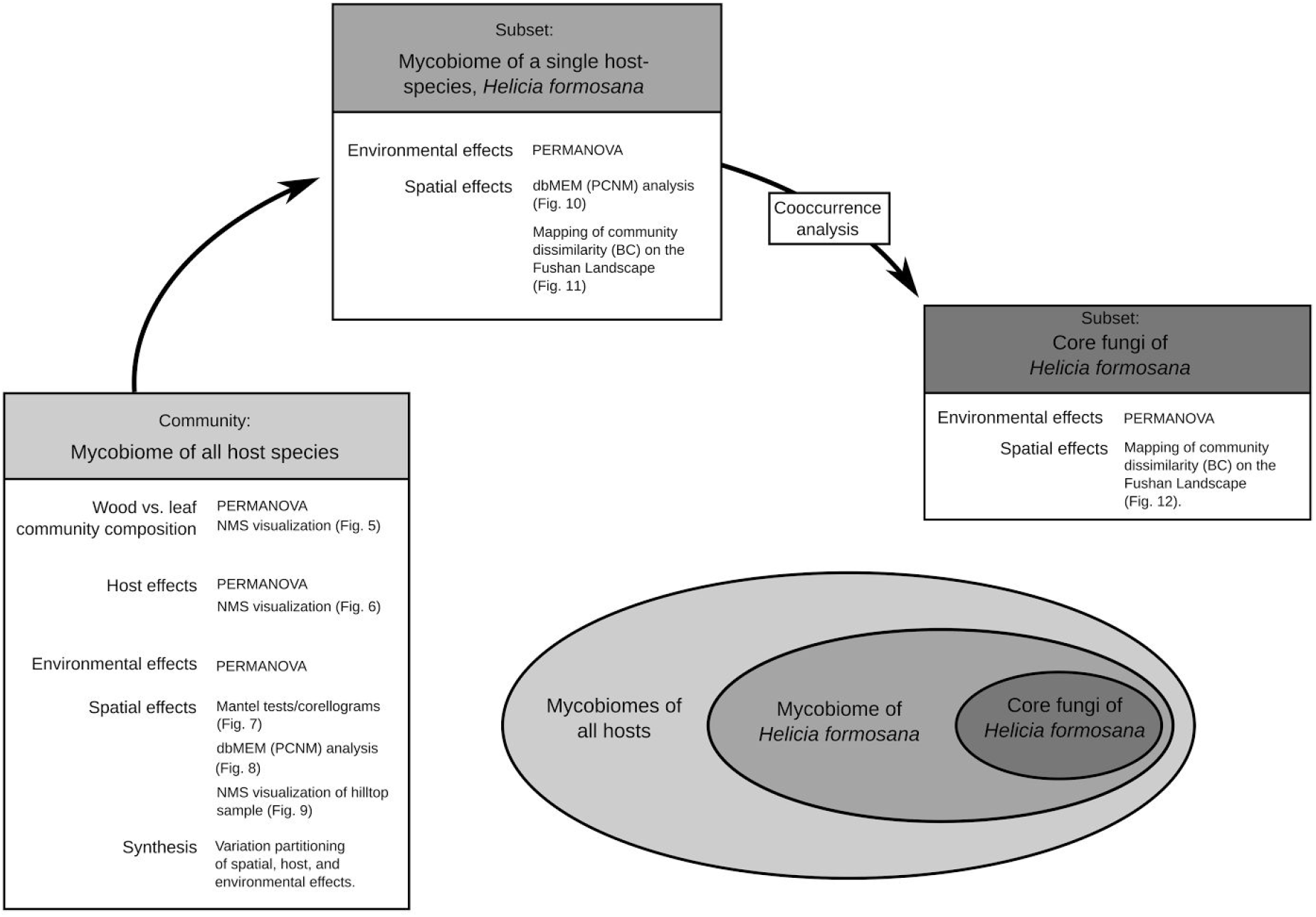
An overview of statistical methods. Analyses begin with broadscale ecological patterns of all wood and leaf samples, then subset to a single host tree species *H. formosana*, and lastly to the patterns of members of core mycobiome of *H. formosana* as defined by cooccurrence patterns. Click here for a higher resolution image.

### Mycobiome of all hosts

Dissimilarity of leaf and wood endophyte communities were modeled and visualized using non-parametric multivariate analysis of variance (NPMANOVA or PERMANOVA) (Anderson 2001), and non-metric multidimensional scaling (NMS). Comparisons between leaf and wood libraries were constrained to only shared OTUs, those that were detected at least once in both leaf and wood tissue, to reduce bias from separate library preparations. Following this, all analyses were for wood and leaf endophyes were conducted separately, in parallel. Effects of host and environmental variables of vegetative community and topography (Fig. 1) on endophyte communities were also modeled individually using PERMANOVA, and results were visualized with NMS when significant.

Spatial trends in endophyte communities were first explored using Mantel tests (Mantel 1967, Legendre 1989) of community dissimilarity matrices against physical distance matrices, and visualized with Mantel multivariate correlograms. For greater resolution of spatial trends, distance-based Moran’s eigenvector maps analysis, also known as Principal Components of Neighbor Matrices (PCNM) analysis, was conducted on our sampling scheme. Following the general statistical pipeline recommended by Legendre et al. (Borcard 2011, Legendre 2012), endophyte community matrices were Hellinger-transformed (Legendre 2001), and “regressed” using Redundancy analysis (RDA) (Legendre 2012, Buttigieg 2014) against all eigenvecters (“PCNM vectors”) resulting from dbMEM analysis. Stepwise model selection was then used to filter the biologically important eigenvectors (Oksanen 2017). The remaining eigenvectors were then inspected visually, and used as independent variables in linear-like models of variation partitioning (see below). Ecological patterns of interest detected in spatial analysis were also visualized by mapping Bray-Curtis distance of all wood or leaf samples from a single point of interest (indicated by PCNM vectors), in NMS ordinations.

Overall patterns of dissimilarity among in our endophyte communities were examined using variation partitioning (Peres-neto 2006, Borcard 2011, Gavilanez 2012, Buttigieg 2014). Variation partitioning attempts to explain patterns of dissimilarity among rows of a response matrix among several explanatory matrices, through comparisons of RDA (or other direct-gradient analysis) models created from all possible combinations of explanatory matrices. Here relative effects of host, environmental, and spatial variables on wood and leaf communities were tested as predictors of endophyte community dissimilarity.

### Mycobiome of a single host, Helicia formosana

To examine ecological patterns of mycobiomes without variation resulting from host tree species, the fungal endophytes of a single host tree, *Helicia formosana* Lour. & Hemsl, were examined. This was the host tree for which the most samples (leaves, n=31; wood n=22) were available. Environmental effects on endophyte community were tested with PERMANOVA models of *H. formosana* wood and leaf endophytes against the environmental variables of vegetation class and topography. Spatial patterns were tested by constructing biologically informative PCNM vectors as above, using the subsetted matrix of sites where samples were from *H. formosana* trees. To further visualize, a single sample of interest indicated by the PCNM vectors was used as a center of comparison for all other samples. Bray-Curtis dissimilarity values resulting from comparison were then plotted onto a map of Fushan FDP.

### Core fungi of Helicia formosana

To test for the presence of a core mycobiome, cooccurrence analysis was conducted on the all-host, all-endophyte species-using a pairwise, probabilistic model (Veech 2013). Core mycobiomes of hosts were defined as the subset of fungi that showed strong cooccurrence associations with a host. Strong associations were defined as those with probabilities under null models of random association corrected to a Benjamini-Hochberg false discovery rate (FDR) of 0.05 or less. Focusing on one host, the results were a species composition matrix of just these core species as columns, with rows of just sites where *H. formosana* was sampled.

Patterns of this subset of core fungi were visualized by first calculating Bray-Curtis dissimilarity distance of each sample (row) of this subsetted “core matrix” from an idealized core mycobiome row that contained all members of the core fungi. These values were then mapped on the Fushan FDP plot.

## Results

### Mycobiome of all hosts

#### Endophyte community composition, wood vs. leaves

After variance-stabilization, the wood endophyte library contained 1477 OTUs and the leaf library contained 794 OTUs. They shared 220 mutually-detected OTUs. (Fig. 4) Both leaf and wood samples were dominated by Ascomycota (91% of OTUs in leaves, 83% in wood), but a larger percentage of wood OTUs matched to Basidiomycota (15% of OTUs in wood, compared to 8% of reads in leaves). This larger percentage of Basidiomycetes was due mostly to a larger diversity of Agaricomycetes and Tremellomycetes present in the wood (Fig. 4). Within Ascomycota, both leaf and wood samples contained high percentages of Sordariomycetes, Dothideomycetes, and Eurotiomycetes. Dothideomycetes were present in higher relative diversity in the wood (32% of all OTUs) than in leaf samples (23% of all OTUs). The opposite was true for Sordariomycetes, which were 41% of leaf endophyte OTUs, compared to 18% of wood OTUs. As noted above, all ecological analyses were transformed to incidence data, so that the basic ecological unit for all following analyses was an non-zero observation of an OTU in a sample after cutoffs were subtracted, regardless of read abundance. Trends in numbers of observations parallel patterns in OTU diversity (Fig. 4); if a class of fungi contained a large diversity of OTUs, it also tended to be observed often throughout the study site.

**Figure 4.**
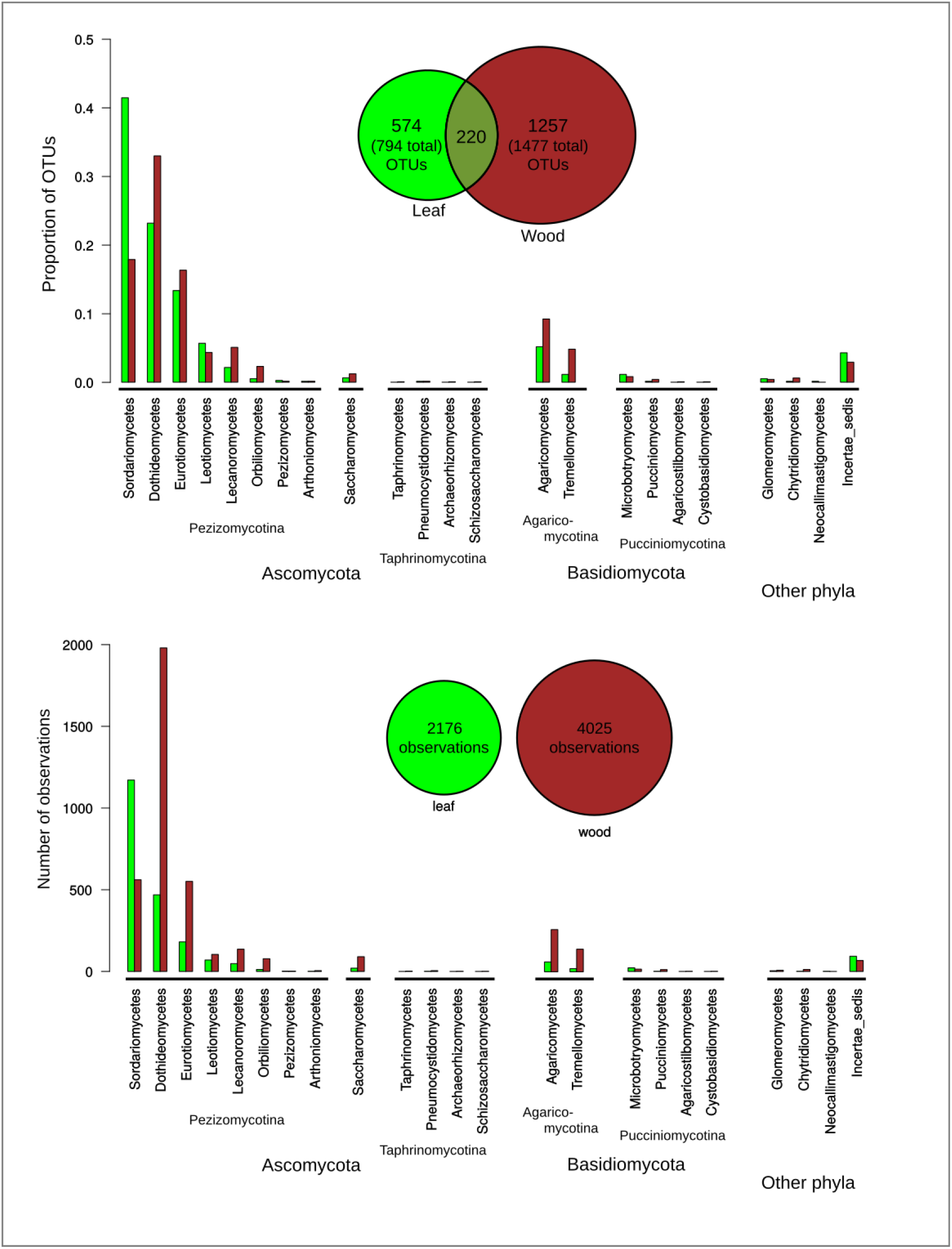
Overview of taxonomic composition of wood and leaf libraries. Top: total numbers of unique OTUs described for each class of Fungi. Bottom: total number of observations of each class. Observations, or presence of a fungal OTU in a sample regardless of read abundance, were the unit of interest for all following analyses, rather than read abundances. Click here for a higher resolution image.

Leaf and wood endophyte communities are distinct, even when analyses are constrained to only species present in both Illumina libraries (PERMANOVA, F(1, 206) = 34.5, p < 0.01, R2 = 0.14, permutations = 10000) (Fig. 5).

**Figure 5.**
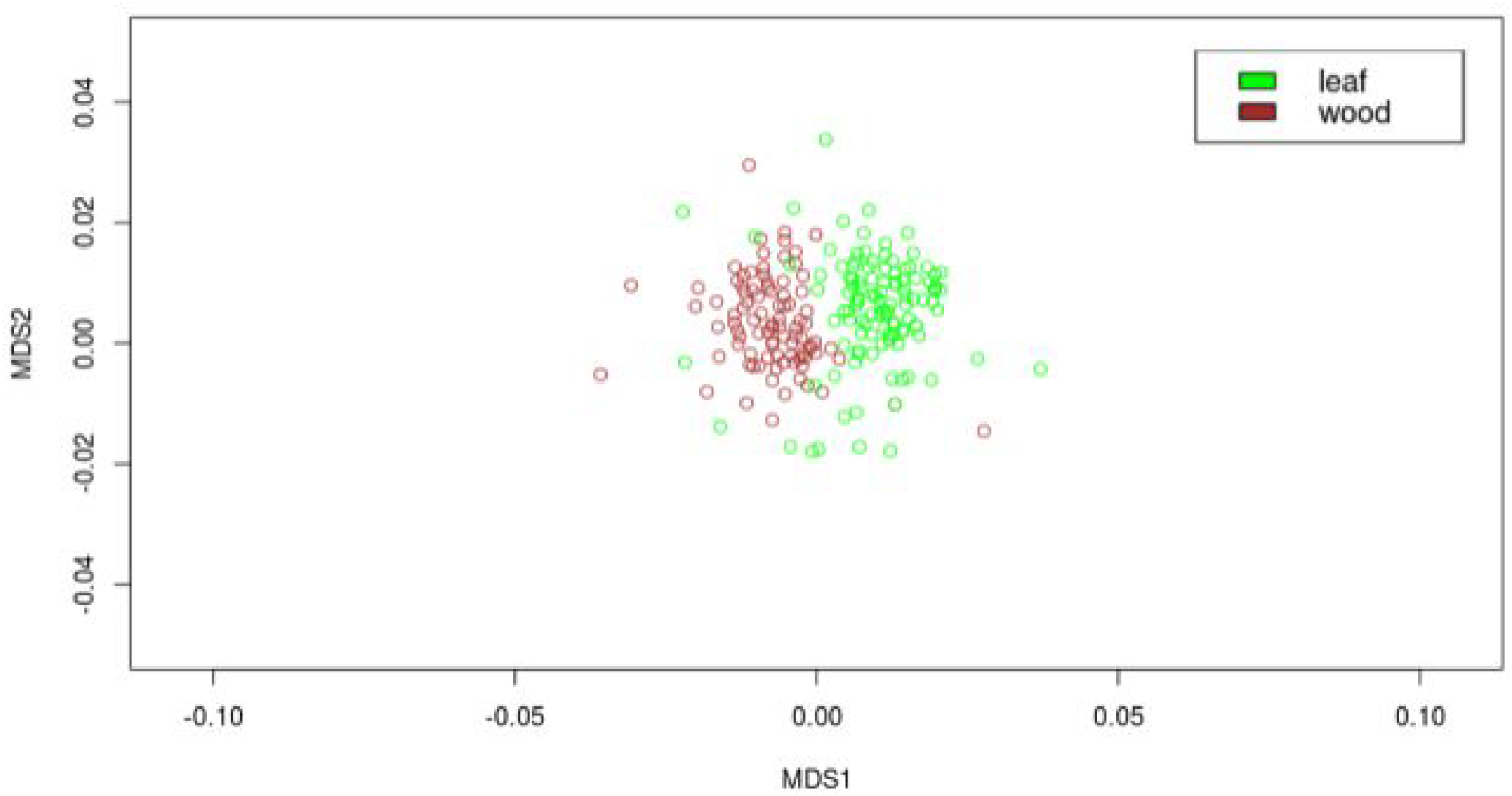
Non-metric multidimensional scaling diagram, comparing leaf and wood endophytes of all host trees, using shared species only. Plot has been scaled in to maximize visibility, two far outliers have been removed. To see entire NMS with outliers, and for a higher resolution image, click here.

#### Host effects on endophyte community composition

Host species is the strongest single predictor of similarity within both leaf (PERMANOVA, F(33, 89) = 2.1, p < 0.01, R^2^ =0.44, permutations = 10000) C and wood endophyte communities (PERMANOVA, F(29,61) = 1.48, p < 0.01, R^2^ = 0.41, permutations = 10000) (Fig. 6).

**Figure 6.**
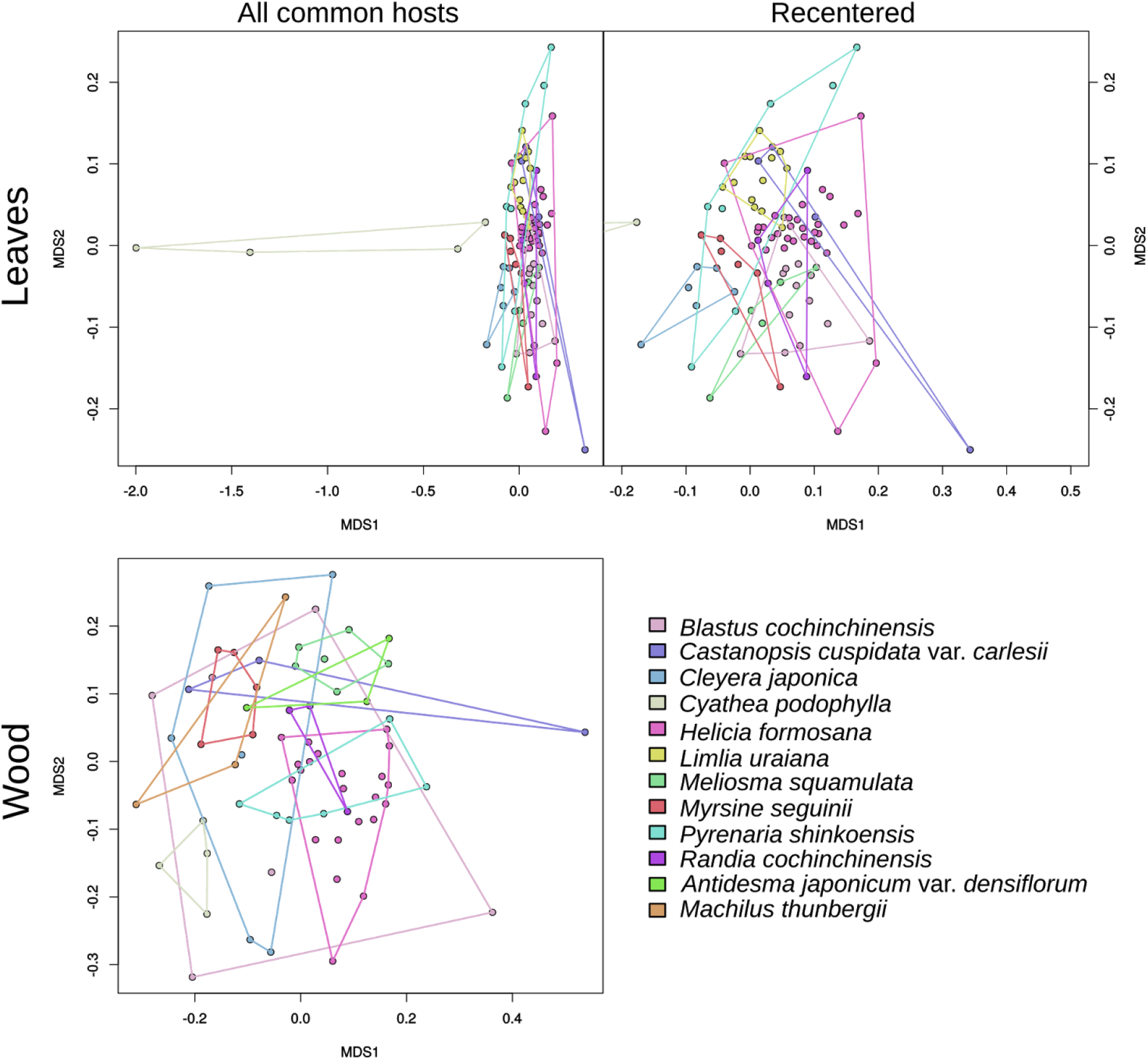
Non-metric multidimensional scaling diagram of endophyte communities, with all tree hosts that were sampled at least 3 times. Leaf plot has been recentered to maximize visibility in upper right, excluding the very unique communities of *Cythea japonica*. Click here for a higher resolution image.

#### Environmental effects on endophyte community composition

Taken alone, composite environmental variables are predictors of similarity in both wood endophyte communities (surrounding above-ground vegetative community: (PERMANOVA, F(3,87) = 1.5, p < 0.01, R^2^ = 0.05, permutations = 10000), micro-topographic conditions (PERMANOVA, F(6,84) = 1.28, p < 0.01, R^2^ = 0.08, permutations = 10000), and also in leaf endophyte communities (surrounding above-ground vegetative community: (PERMANOVA, F(3, 119) = 2.19, p < 0.01, R^2^ = .05, permutations = 10000), micro-topographic conditions (PERMANOVA, F(6, 116) = 1.31, p < 0.01, R^2^ = 0.06, permutations = 10000).

#### Spatial patterns all-host mycobiomes

#### Mantel tests

Wood endophyte community displayed a weak pattern of community-turnover/distance-decay over the entire study area (Mantel's r = 0.07, p = 0.031) (Fig. 7). Leaf communities displayed no global distance decay relationship (Mantel's r = -0.01, p = 0.67), but displayed local negative autocorrelation in comparisons of samples approximately 200 meters apart (Mantel correlogram, Mantel's r = -0.10, p < 0.05) (Fig. 7 indicating that some portion of these samples at this distance apart contained communities more similar than expected under a null model of complete spatial randomness.

**Figure 7.**
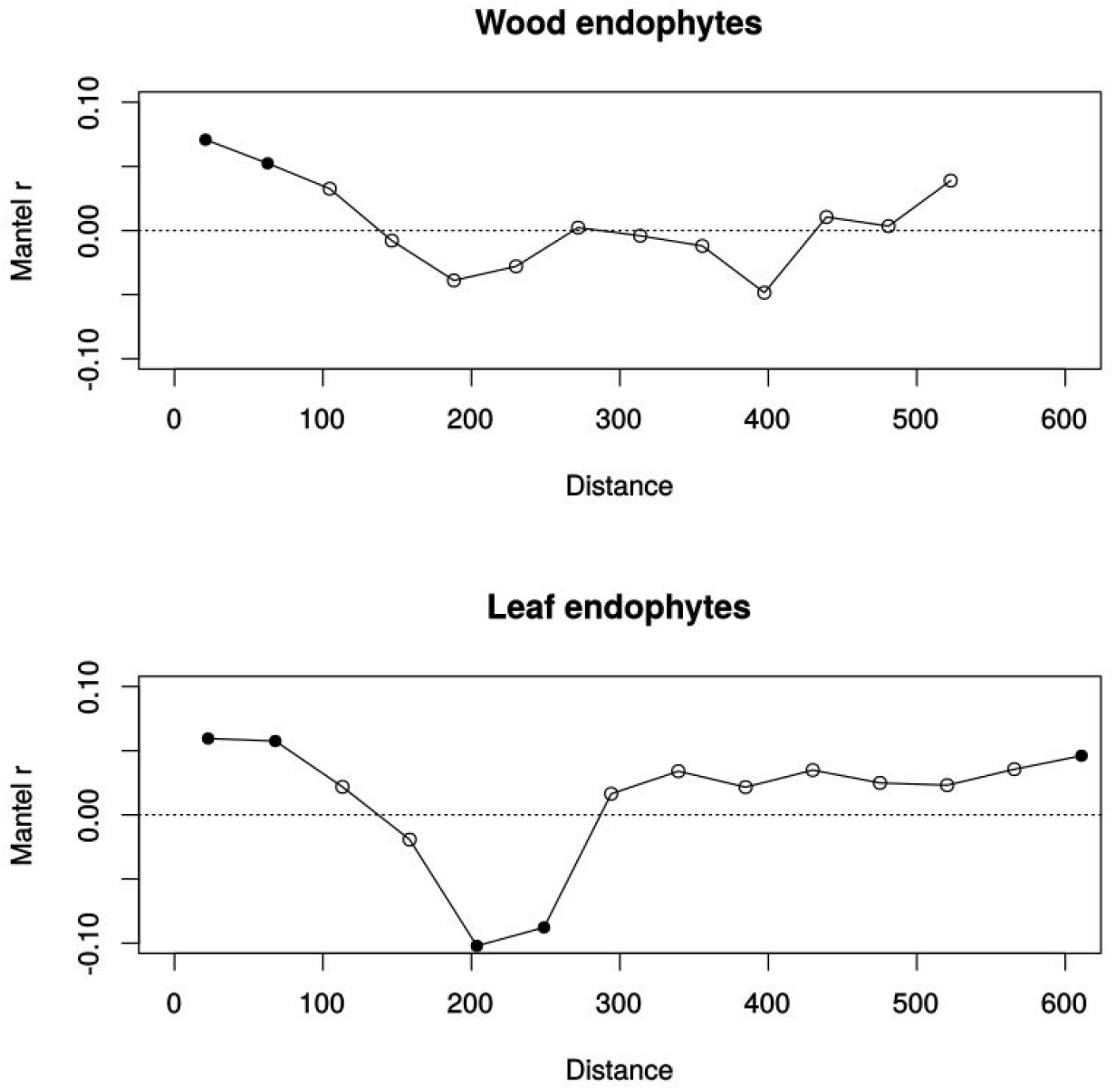
Mantel correlograms of spatial correlation of community dissimilarity of endophyte community. Distance units are meters. Black dots indicate statistical significance. Wood endophytes show weak global distance decay trends. Leaf endophytes do not display global distance decay but have a strong signal of local negative autocorrelation at comparisons around 200 m. Click here for a higher resolution image.

#### dbMEM analyses

Our sampling scheme yielded 5 biologically significant PCNM vectors for leaf samples, explaining 6.6% of endophyte community variation (Redundancy analysis, constrained inertia = 0.06, Unconstrained inertia = 0.89, F(5,117) = 1.65, P < 0.01). Three of five of these PCNM vectors can be considered part of a general north-south pattern that can be combined/detrended as such, and the smallest scale PCNM is probably indicative of endogenous autocorrelation (Borcard 2011). The remaining PCNM vector centers on the hill of the Fushan FDP (Fig. 8), and correlates strongly with environmental variables of topography and vegetative community (Linear model/multiple regression, adj-R^2^=0.64, F(9,113)=25.65, p < 0.01), highlighting this point as important focal point for further comparisons. For leaves, this hilltop point is consistently central in all stable NMS solutions of similarity among all-host comparisons (Fig. 9), and community dissimilarity from this hilltop point is a predictor of dissimilarity among all points ((PERMANOVA, F(1,121) = 8.6, p < 0.01, R^2^ = 0.067, permutations = 10000).

From wood endophyte samples, 4 biologically significant PCNM vectors were described, explaining 6% of variation (Redundancy analysis, constrained inertia = 0.06, Unconstrained inertia = 0.89, F(5,117) = 1.65, P < 0.01). One PCNM correlates strongly with topographical variables (Linear model/multiple regression, adj-R2=.78, F(9,81)=36.39, p < 0.01) and is also centered on the hilltop (Fig. 8). Two of the remaining PCNMs for wood probably represent fine-scale endogenous autocorrelation and the final is not explained well by available variables or visual inspection.

**Figure 8.**
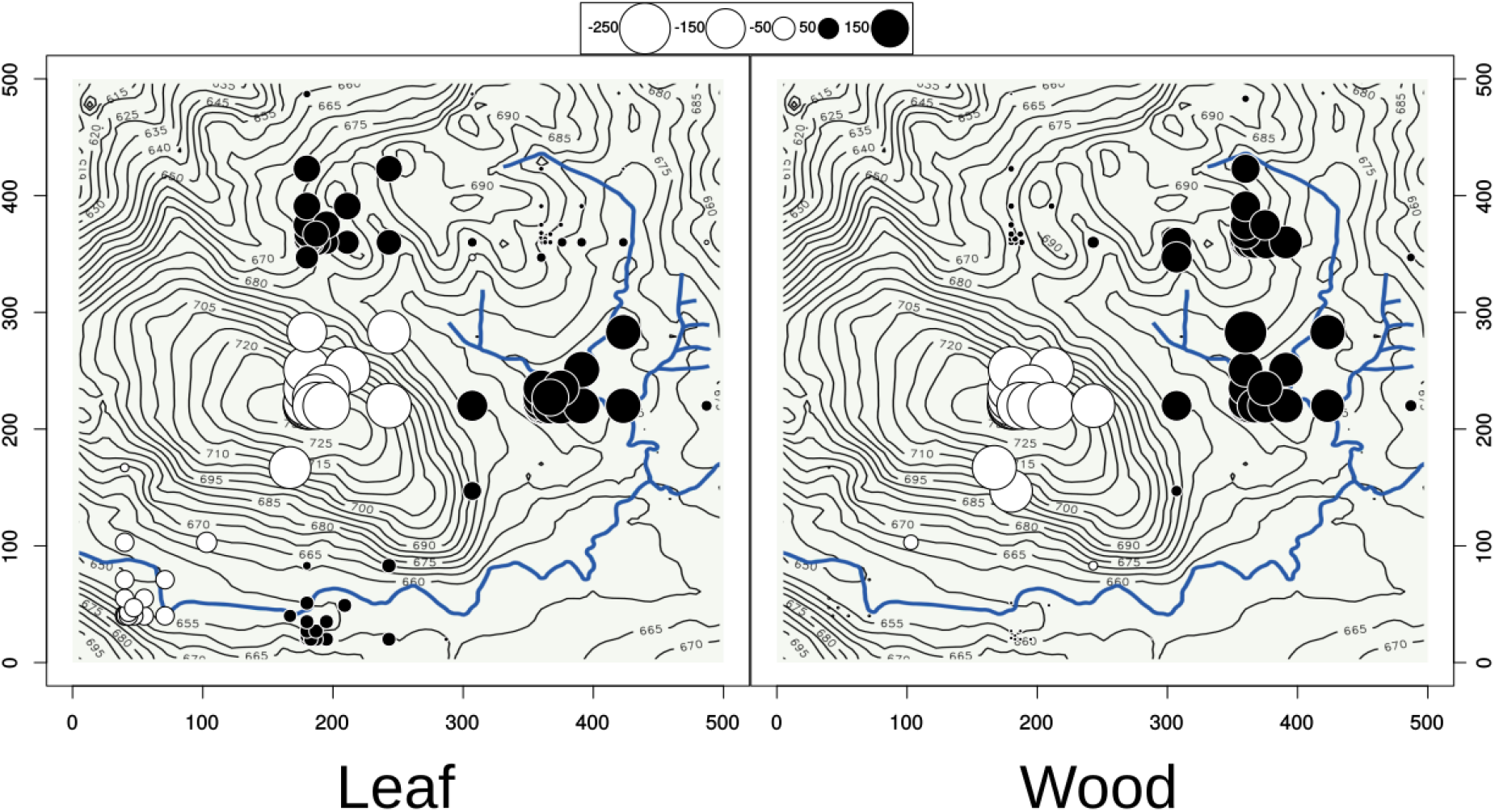
Two PCNM vectors showing patterns of variation of all-host endophyte communities of leaf and wood, plotted over a map of Fushan FDP. Both leaf and wood endophyte communities showed some response to the central hill of the plot. Click here for a higher resolution image.

**Figure 9.**
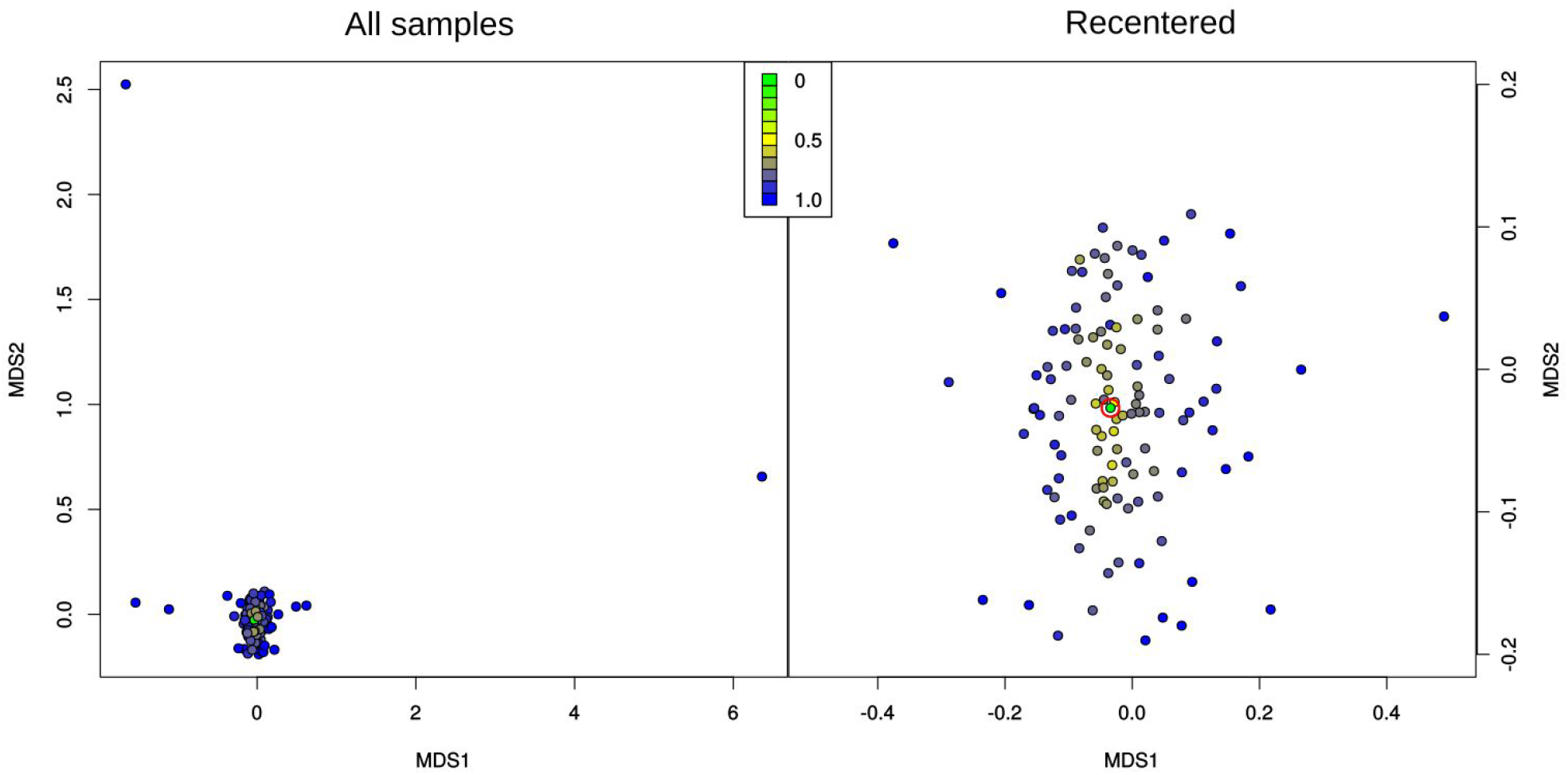
Non-metric multidimensional scaling diagram of leaf endophyte communities. Color indicates community dissimilarity (Bray-Curtis), from a single sample on the central hill of the plot. Dark blue points (BC=1) share no fungal species in common with the hilltop sample, and increase in similarity from yellow to green (BC=0). Leaf plot has been recentered to maximize visibility right, losing 4 samples. Hilltop sample is circled in red on the right. Click here for a higher resolution image.

### Variation partitioning

Most of the variation found among samples in our endophyte communities was unexplained. In wood, host effects explain 5% of total community variation (Redundancy analysis, tested with permutational ANOVA, F(29,54) = 1.20, P = 0.001). Spatial patterns from wood endophytes were not independent of host spatial patterns (Redundancy analysis, tested with permutational ANOVA, F(4,54) = 1.09, P = 0.195). Environmental variables (microtopography and vegetative community) were not observed to explain changes in wood endophyte community directly (0% inertia explained).

Explained variation in leaf endophyte community is also mostly correlated with host effects (10% out of 11% explained; Redundancy analysis, tested with permutational ANOVA, F(9,107) = 2.34, P = 0.001). Independent of host, an additional 1% of leaf endophyte community variation is explained by spatial patterns (Redundancy analysis, tested with permutational ANOVA, F(5,107) = 1.25, P = 0.001). Environmental variables were also not observed to independently explain changes in leaf endophyte community (0% inertia explained).

## Mycobiome of a single host, Helicia formosana

Environmental variables were not found to directly explain any variance in community of *H. formosana* endophytes, for leaves (PERMANOVA, permutations=10000. Topography: F(4,26) = 0.80, p = 0.89, R^2^ = 0.11. Vegetative community: F(3,27) = 1.13, p = 0.24, R^2^ = 0.11), or wood (Topography: F(4,17) = 1.03, p =0.31, R^2^ = 0.20, permutations. Vegetative community: F(3,18) = 1.07, p = 0.23, R^2^ = 0.15). Leaf and wood endophyte community each yielded one biologically significant PCNM vector (RDA, leaves: constrained inertia = 0.044, Unconstrained inertia = 0.72, F(1,29) = 1.78, P < 0.01. RDA, wood: constrained inertia = 0.052, Unconstrained inertia = 0.75, F(1,20) = 1.38, P < 0.01). These PCNMs both display a pattern of dissimilarity centered on the southwest valley (Fig. 10). Centering the Bray-Curtis comparisons on this region shows that leaf samples in this region share fungal OTUs (Fig. 11).

**Figure 10.**
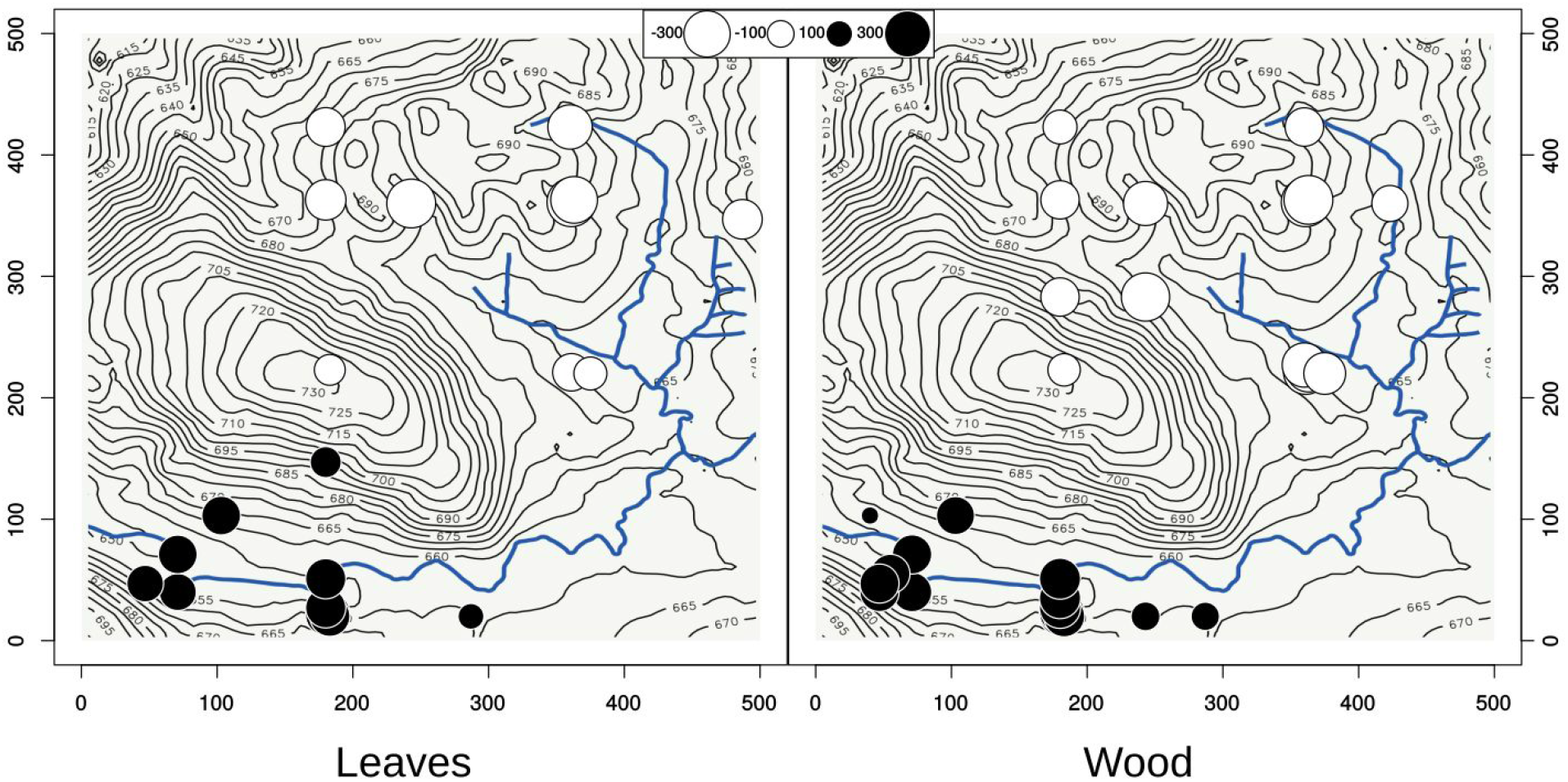
Two PCNM vectors showing patterns of variation of single host-tree, *Helicia formosana*, endophyte communities of leaf and wood, plotted over a map of Fushan FDP. Both leaf and wood endophyte communities display dissimilarity between the plot at large and the southern valley. Click here for a higher resolution image.

**Figure 11.**
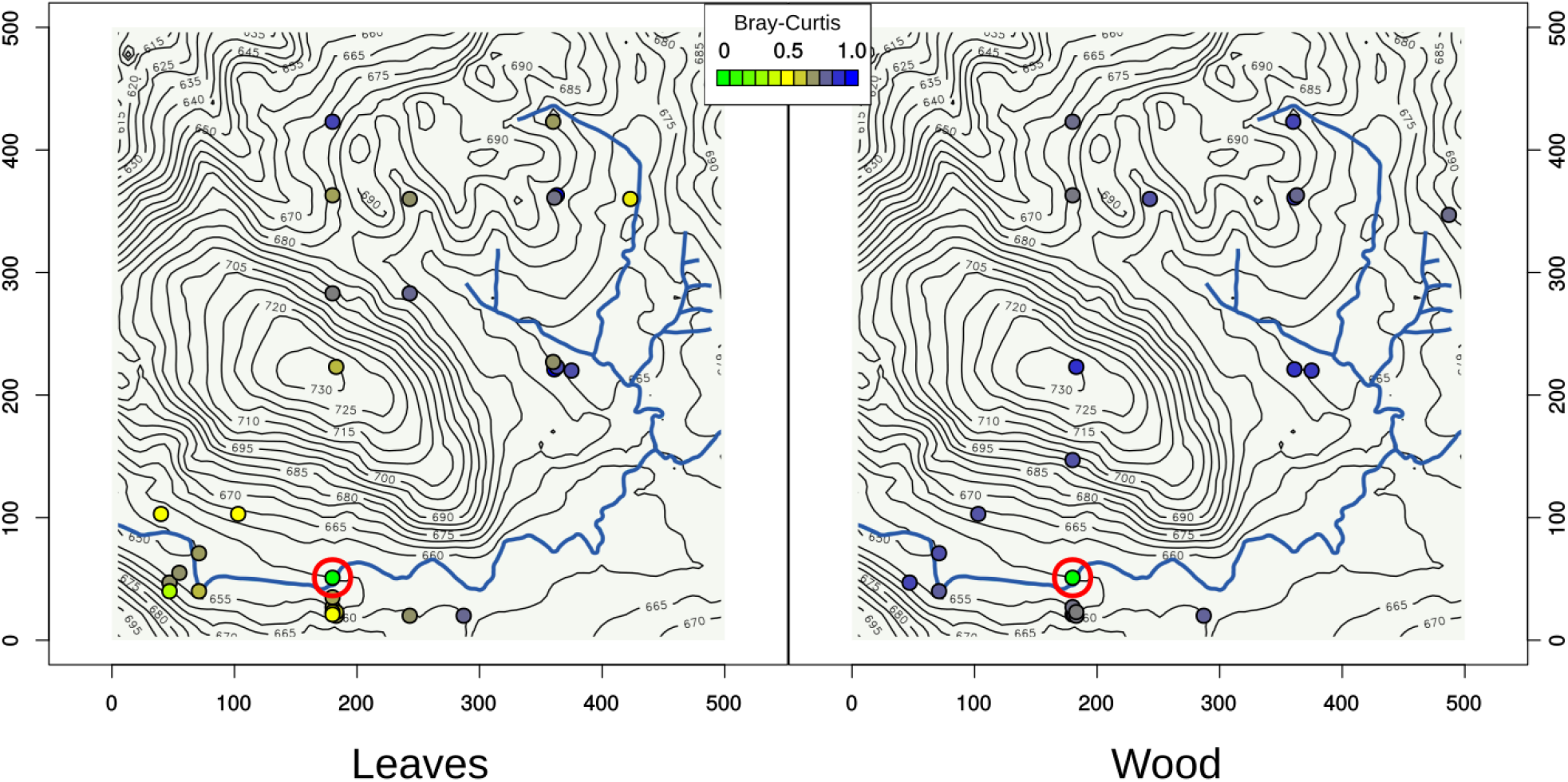
Map of Bray-Curtis dissimilarity values over the Fushan FDP, resulting from comparisons between red circled point all other *Helicia formosana* samples. Dark blue points (BC=1) share no fungal species in common with the circled sample, and increase in similarity from yellow to green (BC=0). Click here for a higher resolution image.

## Cooccurrence analysis

8 out of 774 possible fungal OTUs showed patterns of cooccurrence with *Helicia formosana* in leaf tissue, and 10 out of 1477 possible taxa from wood tissues (Table 1). These fungi were considered members of the *H. formosana* core mycobiome for further analysis.

**Table 1.** Core mycobiome of *Helicia formosana*, defined by cooccurrence patterns. Click here for a higher resolution image.

|  | Taxa | Kingdom | Phylum | Subphylum | Class | Order | Family | Genus |
| --- | --- | --- | --- | --- | --- | --- | --- | --- |
| Leaves | OTU65:128Leaf | Fungi | Ascomycota | Pezizomycotina | Dothideomycetes | Pleosporales |  |  |
|  | OTU75:21Leaf | Fungi | Ascomycota | Pezizomycotina | Sordariomycetes | Diaporthales |  |  |
|  | OTU226:101Leaf | Fungi | Ascomycota | Pezizomycotina | Orbiliomycetes | Orbiliales | Orbiliaceae | Arthrobotrys |
|  | OTU136:130Leaf | Fungi | Basidiomycota | Pucciniomycotina |  |  |  |  |
|  | OTU221:103Leaf | Fungi | Ascomycota | Pezizomycotina | Eurotiomycetes | Eurotiales | Trichocomaceae | Aspergillus |
|  | OTU59:96Leaf | Fungi | Ascomycota | Pezizomycotina | Dothideomycetes | Botryosphaerales | Botryosphaeriaceae | Phyllosticta |
|  | OTU133:116Leaf | Fungi | Zoopagomycota |  |  |  |  |  |
|  | OTU5434:100Leaf | Fungi | Ascomycota | Pezizomycotina | Dothideomycetes | Botryosphaerales | Botryosphaeriaceae | Phyllosticta |
| Wood | OTU284:131Leaf | Fungi | Ascomycota | Pezizomycotina | Eurotiomycetes | Chaetothyriales |  |  |
|  | OTU1126:199wood | Fungi | Ascomycota | Pezizomycotina | Dothideomycetes | Capnodiales |  |  |
|  | OTU256:38Leaf | Fungi | Basidiomycota | Agaricomycotina | Tremellomycetes | Tremellales |  |  |
|  | OTU841:237wood | Fungi | Ascomycota | Pezizomycotina | Dothideomycetes | Capnodiales |  |  |
|  | OTU68:55Leaf | Fungi | Ascomycota | Pezizomycotina | Dothideomycetes |  |  |  |
|  | OTU65:128Leaf | Fungi | Ascomycota | Pezizomycotina | Dothideomycetes | Pleosporales |  |  |
|  | OTU81:81Leaf | Fungi | Ascomycota | Pezizomycotina | Sordariomycetes | Xylariales | Amphisphaeriaceae | Pestalotiopsis |
|  | OTU216:90Leaf | Fungi | Ascomycota | Pezizomycotina | Eurotiomycetes | Verrucariales |  |  |
|  | OTU235:131Leaf | Fungi | Ascomycota | Pezizomycotina | Dothideomycetes | Capnodiales | Capnodiaceae |  |
|  | OTU113:194wood | Fungi | Ascomycota | Pezizomycotina | Eurotiomycetes | Chaetothyriales | Herpotrichiellaceae |  |

## Core fungi of *Helicia formosana*

No direct relationship between topographic and vegetative community and was detected in either leaf endophytes (PERMANOVA, permutations=10000. Topography: F(4,24) = 1.30, p = 0.26, R^2^ = 0.18. Vegetative community: F(3,25) = 0.57, p = 0.79, R^2^ = 0.06), or wood endophytes (PERMANOVA, permutations=10000. Topography: F(4,17) = 1.05, p = 0.35, R^2^ = 0.19. Vegetative community: F(3,18) = 0.86, p = 0.53, R^2^ = 0.13). Visual inspection of spatial patterns show that leaves within the southern valley of the plot contained relatively high proportions of core fungi (Fig. 12). Wood contained high proportions of core fungi consistently throughout the plot (Fig. 12). In leaves, presence or absence of just these core species in *H. formosana* leaf fungal communities is a partial predictor of entire fungal community structure (PERMANOVA, F(1, 29) = 3.38, p < 0.01, R^2^ = .10, permutations = 10000), and for wood endophyte community structure (PERMANOVA, F(1, 20) = 1.29, p = 0.047, R^2^ = 0.06, permutations = 10000).

**Figure 12.**
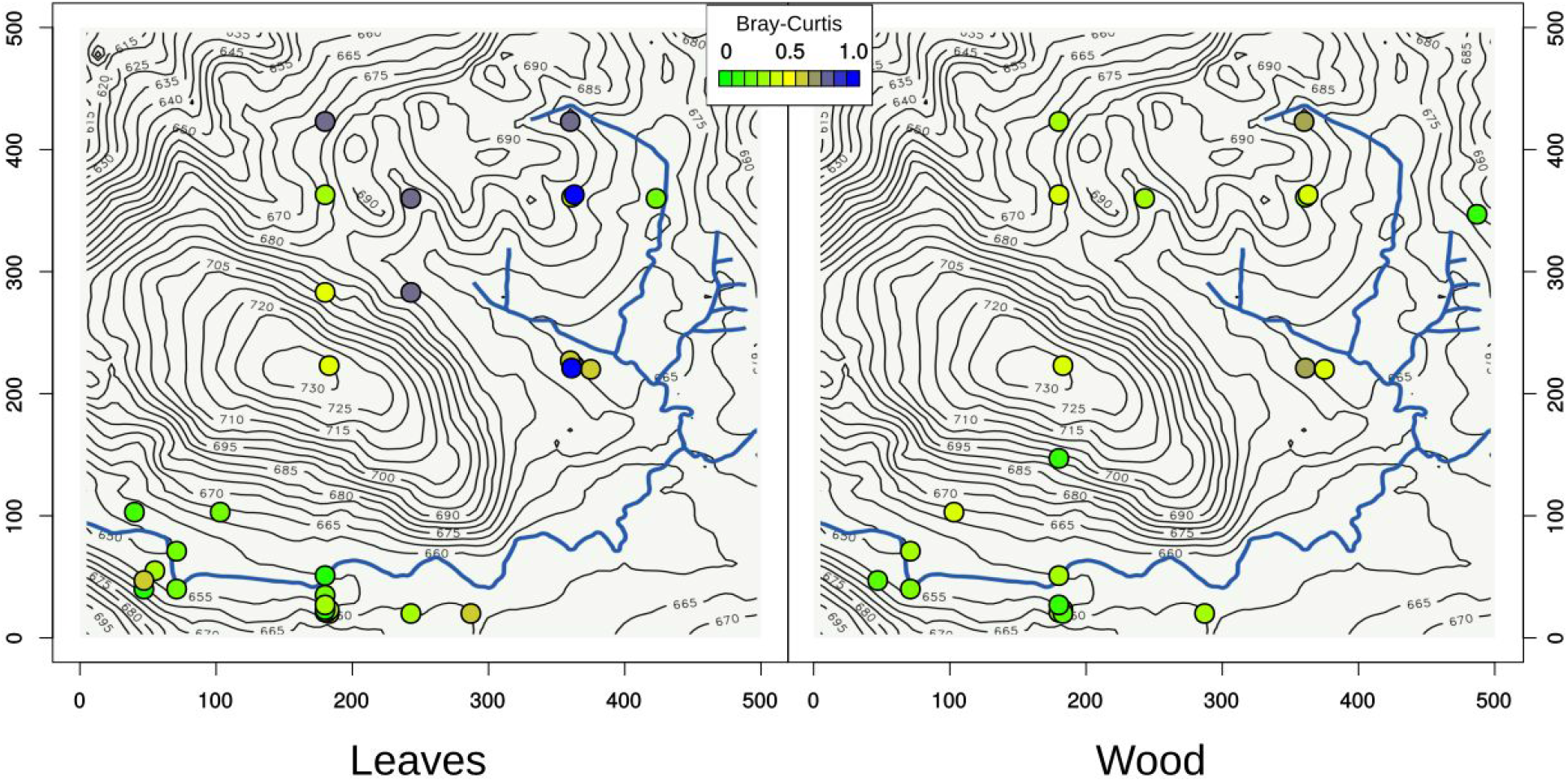
Map of Bray-Curtis dissimilarity values over the Fushan FDP, resulting from comparisons between all *H. formosana* points and the core fungi of the *H. formosana*. Dark blue points (BC=1) contain no species from this set of core fungi, and increase in similarity from yellow to green (BC=0, 100% of core fungi present). Click here for a higher resolution image.

## Summary comparison

The above analysis compared patterns of community dissimilarity at several levels (Fig. 13, Table 2). Wood and leaf endophytes of all host-trees showed an identical, high mean Bray-Curtis dissimilarity among all samples (all-host leaf endophyte mean BC=0.9, sd=0.10; wood endophyte mean BC=0.9, sd=0.07). samples are more similar to one another when considering only one host species, *Helicia formosana* (leaf mean BC=0.78, sd =0.12; wood mean BC = 0.81, sd=0.07). This variation can then be partitioned into two groups: Non-core fungi from these hosts show a similar, high level of dissimilarity among samples (leaf mean BC=0.86, sd =0.11; wood mean BC = 0.86, sd=0.06). As expected, core fungi assemblages from *Helicia* samples have a lower mean BC (leaf mean BC=0.50, sd =0.27; wood mean BC = 0.40, sd=0.17). Leaf core fungi are more dynamic than wood, showing a higher mean Bray-Curtis dissimilarity and greater variance.

**Figure 13.**
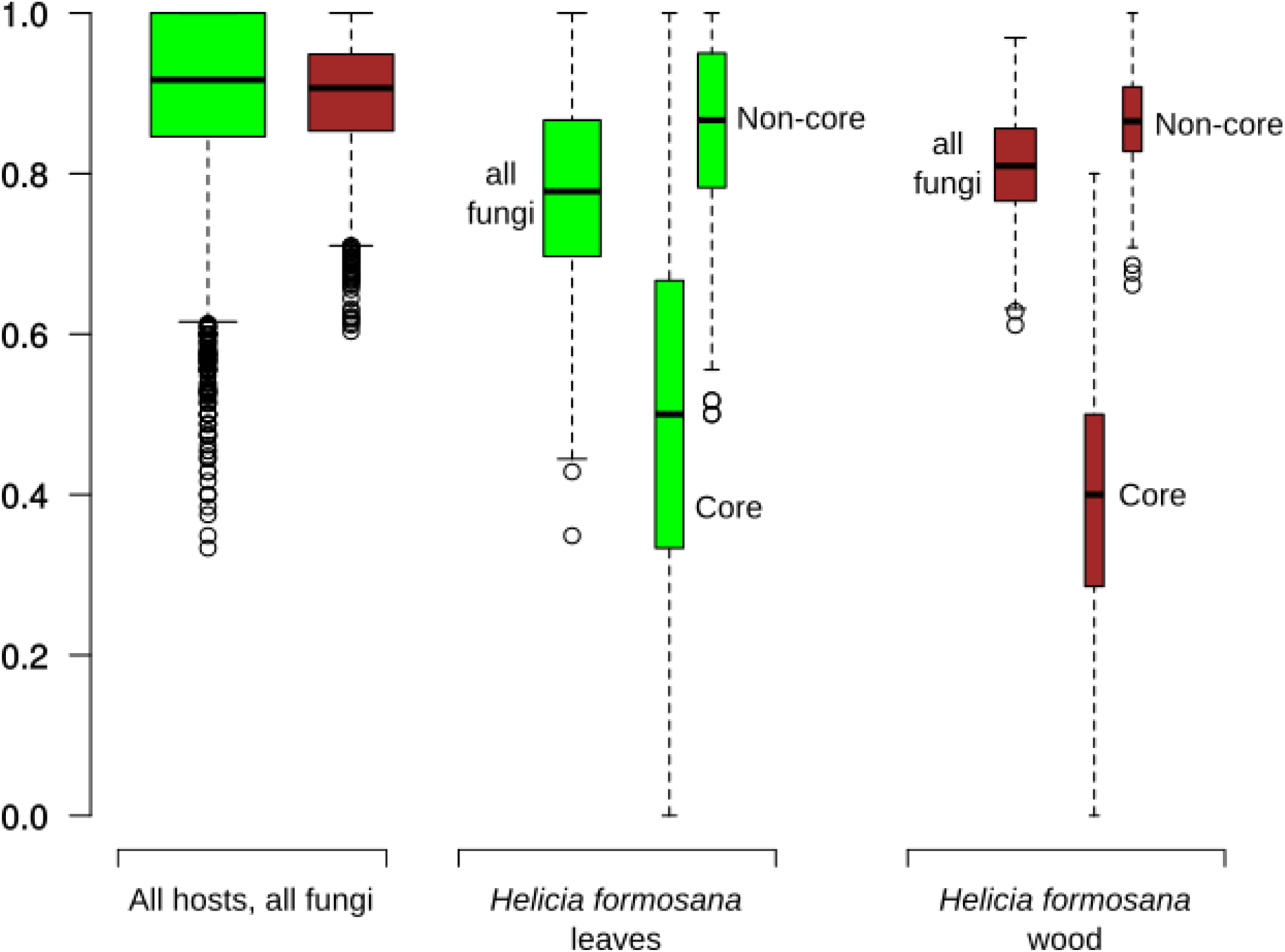
Distribution of Bray-Curtis dissimilarity among sample comparisons of all hosts, and of *Helicia formosana* only. Click here for a higher resolution image.

**Table 2.** Summary mean and standard deviation of Bray-Curtis dissimilarity among sample comparison of all hosts and of *Helicia formosana* only.

| organ | all hosts, all endophytes | *Helicia*, all-endophytes | *Helicia*, non-core | *Helicia*, core-fungi |
| --- | --- | --- | --- | --- |
| leaf | .90(+/- .10) | .78(+/- .12) | .86(+/- .11) | .50(+/- .27) |
| wood | .90(+/- .07) | .81(+/- .07) | .86(+/- .06) | .40(+/- .17) |

## Discussion

The fungal mycobiomes of trees at Fushan FDP are highly variable, and we uncovered only a small part of the reason for their enormous variability. When all host trees are compared, the average dissimilarity between any two trees is extremely high, (Fig. 13, Table 2). Samples become somewhat more similar on average when constrained to a single host, for wood and leaves, a result of the strong effects of host (Fig. 6). But we do see an assemblage of fungi, 8 species in leaves and 10 in wood, “the core” that behave differently. Removing these fungi from consideration brings the mycobiome of their host, *H. formosana*, nearly back to background levels of dissimilarity among samples of the entire study, indicating that these are the species through which host effects are manifested (Fig. 13).

These two sets of core fungi show differing spatial patterns (Fig. 12). In leaves, these core fungi are most consistently present in the southern valley, and are often completely missing in other areas of the study. In wood, they are more “loyal”, and coexist more reliably with *H. formosana* throughout the plot. This may perhaps be due to the high rate of turnover in leaves, which are flushed mostly sterile (Arnold 2003), and are shed within 1 to several years, in contrast with the longer lifespan of woody tissues. Applying terminology proposed by Hamady and Knight (2009), core woody endophytes here may be best described by the “minimal” core model: they are few in number among a large and highly variable microbiome, but are consistently present throughout the study. In contrast, leaf endophytes may be described better by “gradient” or “subpopulation” core models, where a core group of associated microbes may establish with a particular host, but whose presence is highly conditional on space and environment.

Among the endophytes of all hosts, the central hill of FDP was important. We observed in leaves a homogenizing spatial effect with a radius of ~200 m, centered around the hill of the FDP (Fig. 7, 8). The hill of the Fushan plot was central point in the community dissimilarity space of all the samples (Fig. 9). This is surprising, because the hilltop is a very distinct environment from the surrounding lowlands (Fig. 1), which have more in common with each other than with the hilltop. We were limited here by our coarse environmental data in the arguments that can be made for neutral spatial effects versus environmental filters as major predictors. However, this suggests that neutral effects may have been at work: the hilltop may be acting as a dispersal obstacle among the lowland areas, causing local structuring of microbial communities, especially the sheltered southwestern valley, and acting also as a common crossroads through which more widely dispersed microbes must pass. Being the exposed, high point of an area frequently subject to hurricanes, this hilltop may also be a local source of microbial species that are wind-dispersed. Conversely, where we see the most stable cooccurrence relationships are to be found in the relatively sheltered southwestern valley of the FDP.

The presence of a core group of microbes in a host can be seen as a kind of stabilization or structuring of a portion of a host’s microbiome, as a result of interactions among hosts and select microbes. Extensive dispersal and disturbance can disrupt the effects of species interactions and beta diversity/local structure in communities and gene pools (Wright 1940, Cadotte 2006, Vellend 2010). We see that a single, relatively small land feature, a hill representing an 80m elevation gain, can have great effect on the microbes of a landscape, disrupting seemingly strong microbe-host affinities. However, when defining core microbiomes, it may be important to consider the different organs of hosts as very different refugia for microbes: here the more stable environment of woody tissues appeared to host a more consistent assemblage of fungi. Similarly, the leaves *Helicia formosana* trees in the more sheltered southwestern valley held more consistent microbial communities than in more exposed areas of the plot. We conclude that even the strongest biological interactions between microbe and host can be disrupted by neutral processes or environmental changes. This implies that for a consistent core microbiome to develop, either local habitat or host must provide some measure of stability through time and space for local community structuring of microbes to occur.

